# Structural variation turnovers and defective genomes: key drivers for the in vitro evolution of the large double-stranded DNA koi herpesvirus (KHV)

**DOI:** 10.1101/2022.03.10.483410

**Authors:** Nurul Novelia Fuandila, Anne-Sophie Gosselin-Grenet, Marie-Ka Tilak, Sven M Bergmann, Jean-Michel Escoubas, Sandro Klafack, Angela Mariana Lusiastuti, Munti Yuhana, Anna-Sophie Fiston-Lavier, Jean-Christophe Avarre, Emira Cherif

**Affiliations:** ISEM, Univ Montpellier, CNRS, IRD, Montpellier, France; DGIMI, Univ Montpellier, INRAE, Montpellier, France; Institute of Infectology, Friedrich-Loeffer-Institut, Federal Research Institute for Animal Health, Greifswald-Insel Riems, Germany; IHPE, Univ Montpellier, CNRS, Ifremer, UPVD, Montpellier, France; Department of Experimental Animal Facilities and Biorisk Management, Friedrich-Loeffer-Institut, Federal Research Institute for Animal Health, Greifswald-Insel Riems, Germany; Fish Health Laboratory, Research Institute for Freshwater Aquaculture and Fisheries Extension, Bogor, Indonesia; Faculty of Fisheries and Marine Sciences, Bogor Agricultural University, Indonesia; Institut Universitaire de France (IUF)

**Keywords:** KHV, virus evolution, virulence, virus attenuation, structural variations, defective genome, carp

## Abstract

Structural variations (SVs) constitute a significant source of genetic variability in virus genomes. Yet knowledge about SV variability and contribution to the evolutionary process in large double-stranded (ds)DNA viruses is limited. Cyprinid herpesvirus 3 (CyHV-3), also commonly known as koi herpesvirus (KHV), has the largest dsDNA genome within herpesviruses. This virus has become one of the biggest threats to common carp and koi farming, resulting in high morbidity and mortalities of fishes, serious environmental damage, and severe economic losses. A previous study analyzing CyHV-3 virulence evolution during serial passages onto carp cell cultures suggested that CyHV-3 evolves, at least *in vitro*, through an assembly of haplotypes that alternatively become dominant or under-represented. The present study investigates the SV diversity and dynamics in CyHV-3 genome during 99 serial passages in cell culture using, for the first time, ultra-deep whole-genome and amplicon-based sequencing. The results indicate that KHV polymorphism mostly involves SVs. These SVs display a wide distribution along the genome and exhibit high turnover dynamics with a clear bias towards inversion and deletion events. Analysis of the pathogenesis-associated ORF150 region in ten intermediate cell passages highlighted mainly deletion, inversion and insertion variations that deeply altered the structure of ORF150. Our findings indicate that SV turnovers and defective genomes represent key drivers in the viral population dynamics and in vitro evolution of KHV. Thus, the present study can contribute to the basic research needed to design safe live-attenuated vaccines, classically obtained by viral attenuation after serial passages in cell culture.

## Introduction

Viruses have a remarkable ability to adapt to the complex and hostile host immune and physiological constraints. Such capability is directly associated with the viral population’s genetic diversity, and deep characterization of this diversity is the cornerstone of our understanding of virus evolutive response to a new cellular environment. The starkly evident examples are RNA viruses. These viruses, such as HIV, hepatitis C, and influenza, display high mutation rates which generate significant polymorphism levels, allowing the viral population to quickly adapt to newly infected cellular environments and evolve strategies against vaccines and antiviral drugs (Lauring and Andino 2010; Loiseau et al., 2020). However, the genetic diversity has been thoroughly characterized in only a handful of viruses, mainly targeting SNPs (Single Nucleotide Polymorphisms) in RNA viruses (Sanjuán and Domingo-Calap 2016). Although the mutation rate of large double-stranded (ds) DNA viruses is up to four folds lower than that of RNA viruses, due to the use of high-fidelity proofreading polymerases, and most SNPs in dsDNA viruses are neutral and at low frequency, SNP-based approaches were chosen to analyze the genetic diversity in Human cytomegalovirus (HCMV), Autographa californica multiple nucleopolyhedrovirus (AcMNPV) and herpes simplex virus 2 (HSV-2), for example (Renzette et al., 2015; Chateigner et al., 2015; Akhtar et al., 2019).

Structural variations (SVs) play a key role in viral evolutionary processes. Genome rearrangements such as deletions, insertions, duplications and inversions can lead to defective viral genomes (DVGs) (O’Hara et al., 1984; Molenkamp et al., 2000; Vignuzzi and López 2019). Preben von Magnus first identified DVGs in the late 40’s as incomplete influenza viruses that can interfere with the wild-type virus replication (Vignuzzi and López 2019). Since then, the role of DVGs in antiviral immunity, viral persistence and their negative impact on virus replication and production has been established (Bull et al., 2003; Li et al., 2011; Vignuzzi and López 2019; Loiseau et al., 2020). Nowadays, DVGs have been described in most RNA viruses and to a lesser extent in dsDNA viruses (Vignuzzi and López 2019; Loiseau et al., 2020). Despite the critical role of SVs in virus infection dynamics, the knowledge about structural variation diversity, and their evolutionary impact in viral populations, especially those with large dsDNA, is limited.

The large dsDNA Cyprinid herpesvirus 3 (CyHV-3), more commonly known as koi herpesvirus (KHV), is one of the most virulent viruses of fish. It is a lethally infectious agent that infects common carp and koi *(Cyprinus carpio)* at all stages of their life (Hedrick et al., 2000; Haenen et al., 2004). KHV infections are usually associated with high morbidities and mortalities (up to 95%), resulting in serious environmental damages and severe economic losses (Sunarto et al., 2011; Rakus et al., 2013). This threatening virus had a rapid worldwide spread due to global fish trade and international ornamental koi exhibitions (Gotesman et al., 2013). Classified within the family *Alloherpesviridae*, genus *Cyprinivirus*, CyHV-3 is the subject of increasing studies and has become the archetype of alloherpesviruses (Boutier et al., 2015). Despite this “status”, only 19 isolates have been entirely sequenced so far (source: NCBI) since the release of the first complete genome sequences in 2007 (Aoki et al., 2007). Such a low number of full genomes impairs large-scale phylogenomic studies (Gao et al., 2018). On the other hand, KHV infections have been shown to be the result of haplotype mixtures, both *in vivo* and *in vitro* (Hammoumi et al, 2016; Klafack et al, 2019). If mixed-haplotype infections probably represent an additional source of diversification for KHV (Renner and Szpara, 2018), they make genomic comparisons more challenging.

KHV has the largest genome among all known herpesviruses, with a size of approximately 295 kb and 156 predicted open reading frames (ORFs) (Aoki et al. 2007). Several studies focusing on the analysis of viral ORFs have shown the implication of some of them in KHV virulence (Boutier et al., 2015; Fuchs et al. 2014). KHV isolates are known to carry mutations in ORFs that are likely to alter gene functions, and these mutations may vary from virus to virus (Gao et al., 2018) and even within viruses (Hammoumi et al., 2016). Aoki et al (2007) hypothesized that virulent KHV would have arisen from a wild-type ancestor by loss of gene function. However, nearly 15 years later, this hypothesis has still not been tested, probably because of the lack of extensive genomic comparisons. SVs may play a key role in this gene function loss, as recently shown by Klafack et al (2019). These authors conducted a comparative study of a cell culture-propagated isolate that suggested that CyHV-3 evolves through an assemblage of haplotypes whose composition changes within cell passages. This study revealed a deletion of 1,363 bp in the ORF150 of the majority of haplotypes after 78 passages (P78), which was not detected after 99 passages. Furthermore, experimental infections showed that the virus passaged 78 times was much less virulent compared to the original wild-type on the one hand and slightly less virulent compared to the same virus passaged 99 times (P99), highlighting the potentially important role of the ORF150 in the virulence of KHV. Besides, this study demonstrated that haplotype assemblies evolve very rapidly along successive *in vitro* cell passages during infectious cycles, and raised many questions regarding the mechanisms leading to such rapid gene loss and gain *in vitro.*

The present study sought to characterize the SV diversity and dynamics in the KHV genome using viruses propagated onto cell cultures. First, P78 and P99 whole virus genomes were sequenced using ultra-deep long-read sequencing, a first with KHV. Then, the obviously pathogenesis-associated ORF150 region (~5 kb) was sequenced in ten intermediate successive cell passages through an Oxford nanopore® amplicon-based sequencing approach to gain insights into the gene loss and gain mechanisms.

## Methods

### Extraction of high molecular weight DNA from P78 and P99 cell culture passages

The virus isolate used in this study was the same as that previously described in Klafack et al. (2019), *i.e.* an isolate collected from an infected koi in Taiwan (KHV-T) and passed 99 times onto common carp brain (CCB) cells. Considering previous results, a special focus was made on passages 78 (P78) and 99 (P99). Genomic DNA was extracted from cell cultures stored at −80°C, using the MagAttract HMW DNA Kit (Qiagen). Each frozen culture was thawed quickly in a 37°C water bath, equilibrated to room temperature (25°C) and divided into 12 cell culture aliquots of 250 μL. Tubes were centrifuged at 3,000× g for 1 minute and supernatants were transferred into new 2-mL tubes containing 200 μL of proteinase K and RNase A solution. DNA was subsequently extracted according to the manufacturer’s recommendations and eluted in 200 μL distilled water provided in the kit. The 12 replicates of each sample were pooled together and evaporated at room temperature using a vacuum concentrator, to reach a final volume of around 60 μL. Concentrated DNA was quantified by fluorometry (Qubit, ThermoFisher Scientific) and its quality was evaluated by spectrophotometry (Nanodrop 2100) and agarose gel electrophoresis. The final concentration of P78 and P99 was 14.4 and 2.6 ng·μL^-1^, respectively.

### Quantitative PCR assays

Quantitative PCR (qPCR) was applied to evaluate cellular and viral DNA ratio. Two sets of primers were used: primers targeting the ORF150 of CyHV-3 (GenBank #AP008984.1, KHV-J, nt 259,965-260,110: 5’-GAGCGAGGAACTCTACACAAC-3’ and 5’-GGTAAGGGTAAAGCAGACCATC-3’) and primers targeting the glucokinase gene of *Cyprinus carpio* (GenBank #AF053332.2, nt 225-293: 5’-ACTGCGAGTGGAGACACAT-3’ and 5’-TCAGGTGTGGAGGGGACAT-3’). Amplification reactions contained 1 μL of 2X SYBR Green I Master mix (Roche), 200 nM of each primer, and 1 μl of template DNA in a final volume of 10 μL. Amplifications were carried out in a LightCycler 480 (Roche) and cycling conditions consisted of an initial denaturation at 95°C for 5 min followed by 45 cycles of amplification at 95°C for 10 sec, annealing at 60°C for 20 sec and elongation at 72°C for 10 sec with a single fluorescence measurement. After amplification, a melting step was applied, which comprised a denaturation at 95°C for 5 sec, a renaturation at 65°C for 60 sec and a heating step from 65 to 97°C with a ramp of 0.1°C per second and a continuous fluorescence acquisition. Specificity of amplification was verified by visual inspection of the melting profiles, and the ratio between cellular and viral DNA was estimated as 2^-ΔCq^, assuming that each primer pair has an amplification efficiency close to 2 and that each amplicon is present as a single copy per genome.

### Genomic library preparation and Oxford Nanopore whole genome sequencing

High-quality genomic DNA from the two samples (P78 and P99) was sequenced using Oxford Nanopore technology®. DNA libraries were prepared using the Ligation Sequencing Kit (SQK-LSK109) according to the manufacturer’s instructions (Oxford Nanopore®). A total input amount of 48 μL (corresponding to 692 ng of P78 and 124 ng of P99 high molecular weight DNA) was used for sequencing library preparation. DNA was first end-repaired using NEBNext FFPE DNA Repair Mix and NEBNext Ultra II End repair, and then cleaned up with Agencourt AMPure XP beads (Beckman Coulter Inc) at a 1:1 bead to DNA ratio. Sixty-one μL of clean-up elution were transferred into a new 1.5-mL tube for subsequent adapter ligation. Adapter ligation was achieved using NEBNext Quick T4 DNA ligase adapter mix (AMX), ligation buffer (LNB), Long Fragment Buffer (LFB) and Elution Buffer (EB), following the provider’s recommendations. The quantity of the retained DNA fragments was measured again by Qubit fluorometry. The final P78 and P99 DNA amounts were 346 ng and 94 ng, respectively. Each library was directly sequenced on a separate R9.4.1 flow cell using a MinION sequencing device. Sequencing runs were controlled with MinKNOW version 0.49.3.7 and operated for about 30 hours.

### Amplicon-based Oxford nanopore sequencing

To specifically investigate ORF150 region, a fragment of ~4.3 kb encompassing the whole ORF150 of CyHV-3 and part of the ORF149 and ORF 151 (nt 257,103-261,345 according to GenBank #AP008984) was sequenced from various intermediate passages of KHV-T (P10, P20, P30, P40, P50, P70, P78, P80, P90 and P99). A ligase-free protocol was used to limit the risk of potential artifacts linked with sample preparation, e.g. the creation of chimeric sequences during the end-repair or ligation steps (White et al., 2017). Genomic DNA was extracted from cell cultures stored at −80°C, using the Nucleospin virus kit (Macherey-Nagel). Purified DNA was subsequently used for PCR amplification with the following primers: 5’-TGGGCGCAATCAAGATGT-3’ (F) and 5’-TGAAGTGTCAGGGTCAGAGT-3’ (R). PCR was performed with GoTaq G2 DNA Polymerase (Promega) in a final volume of 40 μL containing 1 μL of total genomic DNA, 10 μL of Taq Buffer, 5 μL of dNTPs 2 mM, 2.5 μL of MgCl_2_ and 1.25 μL of each primer (10 μM). Cycling conditions were as follows: initial denaturation at 95°C for 10 min, amplification with 40 cycles of 95°C for 10 sec, 60°C for 20 sec, 72°C for 3 min, and final extension at 72°C for 5 min. PCR products were purified using 1X Agencourt AMPure XP beads, tested for purity using the NanoDrop™ One spectrophotometer (ThermoFisher Scientific), and quantified fluorometrically using the Qubit dsDNA High sensitivity kit. DNA libraries were prepared using the Rapid Barcoding kit (SQK-RBK004), following the manufacturer’s instructions. For each sample, 400 ng of purified amplicon were adjusted with nuclease-free water to a total volume of 7.5 μL and supplemented with 2.5 μL of Fragmentation Mix RB01-4 (one for each sample). Four barcoded samples were combined with an equimolar ratio by mixing 2.5 μL of each sample in a total volume of 10 μL. Furthermore, as described above, two additional barcodes RB01-2 were used with the passages P78 and P99. Pooled libraries were sequenced on 3 R9.4.1 flow cells (4 barcodes, 4 barcodes, 2 barcodes) for 24 hours and sequencing runs were controlled with MinKNOW version 0.49.3.7.

### DNA sequence analysis

For each sample, bases from raw FAST5 files with a *pass* sequencing tag were recalled using the high-accuracy model of ONT Guppy basecalling software version 4.0.15 which improves the basecalling accuracy. The obtained fastq files were filtered to keep reads with a length ≥ 2 kb, in order to span large structural variants. After the mapping step using minimap2 (Li, 2018), the sequencing depth was calculated for each sample using the *<u>plotCoverage</u>* tool implemented in deepTools2.0 tool suite (Ramírez *et al.*, 2016). Sequencing coverage was assessed with the bamCoverage tool from the same tool suite and normalized using the RPGC (reads per genome coverage) method (Figure 1).

**Figure 1:**
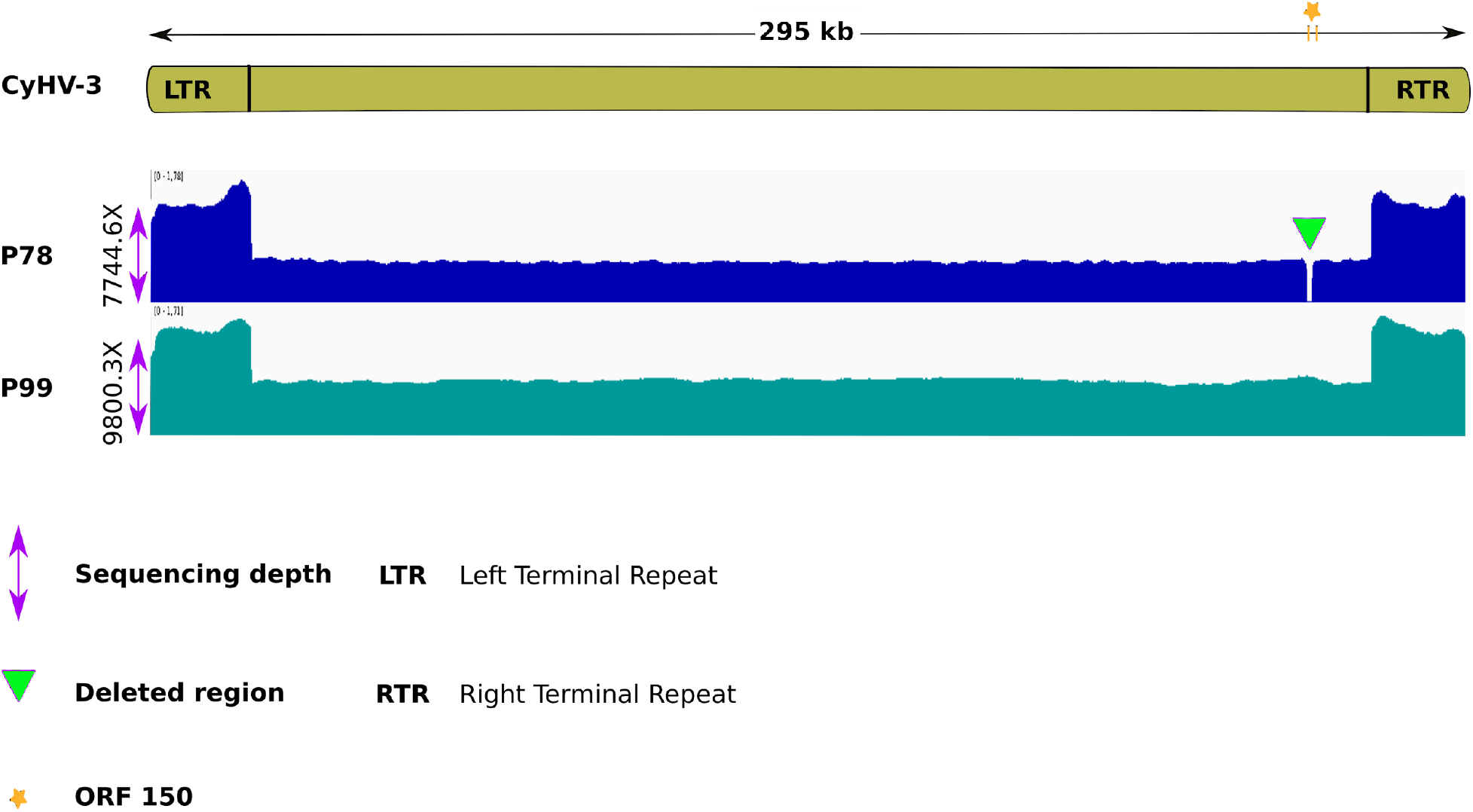
Normalized sequencing coverage for P78 and P99 samples using the RPGC (reads per genome coverage) method. Both P78 and P99 genomes were totally covered by the sequencing. For P78, the coverage break (green triangle) corresponds to the 1.3 kb deletion.

### Structural variant detection

To detect structural variants (SVs) in the P78 and P99 whole genomes, a mapping step followed by BAM filtering was performed. Two aligners were used to map the raw long-reads against the KHV-J AP008984.1 reference genome: minimap2 (Li, 2018) and NGMLR (Sedlazeck et al., 2018). BAM files were then filtered using the option ‘-F’ of Samtools view (Li et al., 2018) with the flag “4” to keep only mapped reads and with the flag “0×800” to remove chimeric reads (inconsistent/supplemental mapping). For P78 and P99, 99.33% and 97.77% of reads were mapped, respectively. Chimeric reads represented 28.05% and 17.74% of the mapped reads in P78 and P99, respectively. The resulting filtered BAM files from each mapper were used as input data for SV caller, Sniffles (Sedlazeck et al., 2018). Only SVs ≥ 30 bp and supported by at least 10 reads were kept in the final VCF files. A cross-validation step was performed using SURVIVOR (Jeffares et al., 2017) by extracting common SVs from each mapper/caller combination for each sample (Figure S1 (Cherif, 2022)). Although the KHV-J reference genome used for the mapping is phylogenetically close to the KHV-T isolate, some genetic diversity exists (Klafack et al. 2017). Hence, a pairwise comparison between P78 and P99 was made to exclude inter-isolate SVs.

The distribution of the different SVs along with P78 and P99 genomes was assessed by estimating their occurrences using a 5 kb sliding window. SNPs and Indels variants were called in P78 and P99 by *medaka_variant* implemented in medaka (1.4.4) using KHV-J AP008984.1 as a reference genome. To detect structural variants in the amplified region (257,103-261,345) of P10 to P99 samples, a size filtering step using *guppyplex* was added to the steps described above. Only reads from 1.5 kb to 8 kb were used for the analysis.

## Results

### Main features of sequencing data for P78 and P99

A total of 4,900,000 and 2,293,830 long-reads were obtained for P78 and P99, respectively. After filtering, 462,982 long-reads with an average length of 4.96 kb were retained for P78, and 418,034 reads with an average length of 7.06 kb for P99 (Table 1, TableS1 (Cherif, 2022)).100% of the sampled bases from the P78 genome had at least 5,000 overlapping reads and 100% of the sampled bases from the P99 genome had at least 7,500 overlapping reads (Figure S2 (Cherif, 2022)). The sequencing data covered both P78 and P99 genomes (Figure 1).

**Table 1.** Main features of reads obtained for each genome after the read length filtering step. min_len= minimum length, avg_len= average length, max_len= maximum length.

| Sample | reads | min_len | avg_len | max_len | N50 | Q20(%) | Q30(%) |
| --- | --- | --- | --- | --- | --- | --- | --- |
| P78 | 462982 | 2000 | 4960.8 | 91494 | 6091 | 59.53 | 17.71 |
| P99 | 418034 | 2000 | 7061.4 | 176512 | 8845 | 58.27 | 19.22 |

N50= the length for which all sequenced reads of that length or greater sum to 50% of the set's total size Q20= 1 in 100 probability of an incorrect base call, Q30= 1 in 1000 probability of an incorrect base call. Here 59.53% of P78 bases=Q20 and 58.27% of P99 bases=Q20.

The ratio between cellular and viral DNA, calculated by qPCR, was 34800 and 33200 for P78 and P99, and the percent of mapped reads 99.33% and 97.77%, respectively.

### SV distribution in P78 and P99

For P78, the mapper/caller combination minimap2/Sniffles detected 731 structural variations (SVs), and the combination NGMLR/Sniffles detected 460 SVs (Table S2 (Cherif, 2022)). For P99, the combination minimap2/Sniffles detected 210 SVs and NGMLR/Sniffles detected 397 SVs (Table S2 (Cherif, 2022)). Independently from the mapper/caller combination that was used, P78 showed more SVs than P99 (Table S2 (Cherif, 2022)). After the cross-validation step (extracting common SVs from each mapper/caller combination), 236 and 87 SVs were kept for P78 and P99, respectively. For comparison, the number of small variations (with a size < 30 nt, which were excluded from this study) amounted to 57 and 77 for P78 and P99, respectively.

In both samples, inversions and deletions were the most prevalent SVs. In P78, inversions represented 75% of the events and deletions 20% (Figure 2.A). In P99, 80% of the SVs were inversions and 11% were deletions (Figure 2.A). Inversions were found along the entire genome, with the highest number detected within the [70-75 kb] window in P78 and within the [235-245 kb] and [270-275 kb] windows in P99 (Figure 2.B). In P78, deletions were mainly detected within [10-40 kb], [70-100 kb] and [245-270 kb] windows (Figure 2.B). In P99, the two major deletions were found within [10-20 kb] and [205-215 kb] regions (Figure 2.B). The frequencies of these SVs were low and did not exceed 1% of the total reads (with a few exceptions, Table S3 (Cherif, 2022)). In spite of such low frequencies, it is interesting to note that the most frequent SVs were located in ORFs potentially involved in DNA replication and encapsidation, e.g. ORF33, 46, 47, 55 (Aoki et al, 2007; Table S3 (Cherif, 2022)).

**Figure 2:**
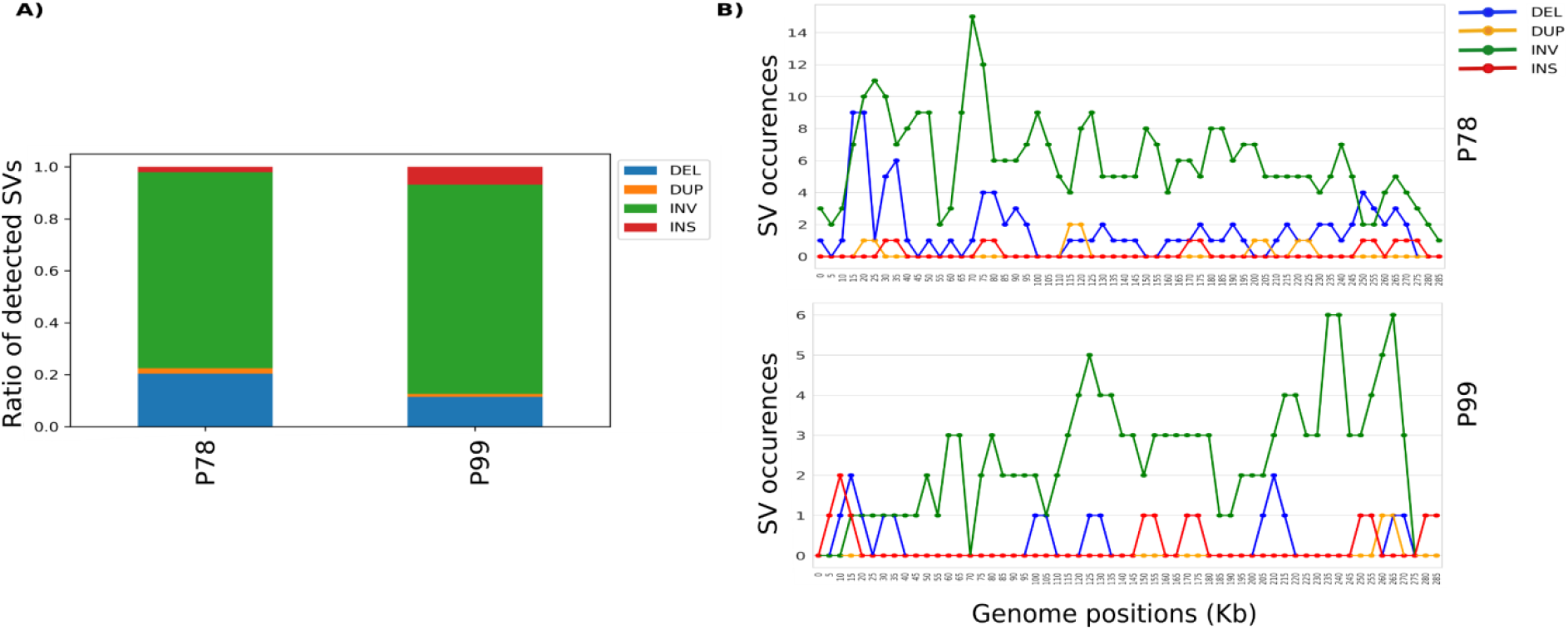
Ratio and distribution of SVs detected in P78 and P99 genomes. **A)** Frequency of each type of SV detected in P78 and P99. **B)** Occurrences and distribution of each SV type in the P78 and P99 genomes using a 5kb window.

Altogether, these results highlight high SV turnover dynamics during the *in vitro* infection cycles (from 78 passages to 99) with a clear trend, or bias, towards inversion and deletion events.

### Dynamics and impacts of SVs in ORF150 region

Taking advantage of the high-resolution SV detection provided by the long-read sequencing, we looked for the SV events around the potential virulence-linked ORF150 in P78 and P99 (nt 257,103-261,345 according to AP008984.1). Results confirmed that P99 had a reference-like profile with an unmodified ORF150. In P78, the deletion (nt 258,154-259,517; D258153) was found in 6,902 reads (100% of the reads), whereas the reference haplotype was also detected in 30 reads, representing 0.44% of the total 6,734 supporting reads (Figure 3, Tables S4 (Cherif, 2022) (Cherif, 2022)). Surprisingly, 26 reads revealed a haplotype as yet unidentified (INV258153), consisting of an inversion of the same length (1,363 bp) and at the same breakpoints as the deletion. The inverted haplotype (INV258153) in P78 deeply altered the ORF150, by inverting the first 1200 bp of the ORF and 160 bp of the 5’UTR in the middle of the ORF (Figure 3).

**Figure 3:**
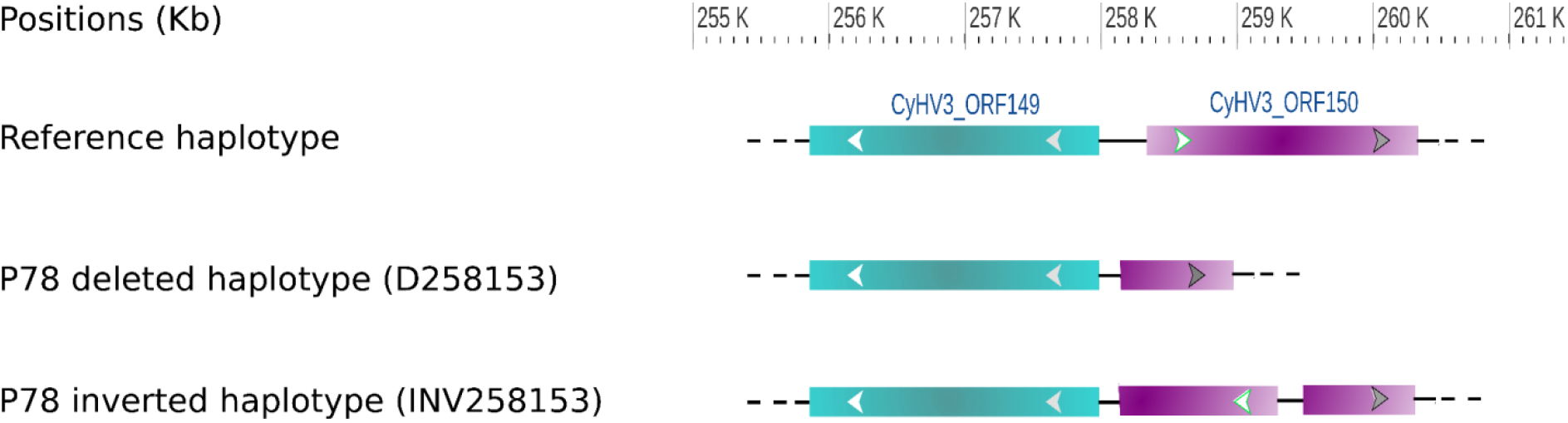
Impact of structural variations D258153 and INV258153 on the P78 genomic structure. The inversion is highlighted by an inverted arrow compared to the reference haplotype.

In order to trace the unexpected dynamics of gain and loss of the full ORF150 along passages, we searched for the SV turnovers during 10 intermediate passages (P10, P20, P30, P40, P50, P70, P78, P80, P90, P99). This analysis revealed the presence of haplotype D258153 at low frequency (from 0.05 to 0.15% of the reads) in passages P10 to P40 and a strong increase in its frequency at P50 (88.7% of the reads) (Figure 4). The frequency of the haplotype D258153 reached a maximum at P78 (100% of the reads) then dropped quickly at P80 (30.7% of the reads) to stabilize at low frequency (0.31% of readings) at P90., as during the first 40 passages (Figure 4, Table S4 (Cherif, 2022)). Interestingly, shorter deletions of 119 and 881 bp were observed near the 5’ end of the ORF150 in P40 and P80, respectively, at low frequencies (0.42 % in P40 and 0.18% in P80) (Figure 4, Table S4 (Cherif, 2022)). The haplotype D258153 completely disappeared at passage 99 (Figure 4, Table S4 (Cherif, 2022)).

**Figure 4:**
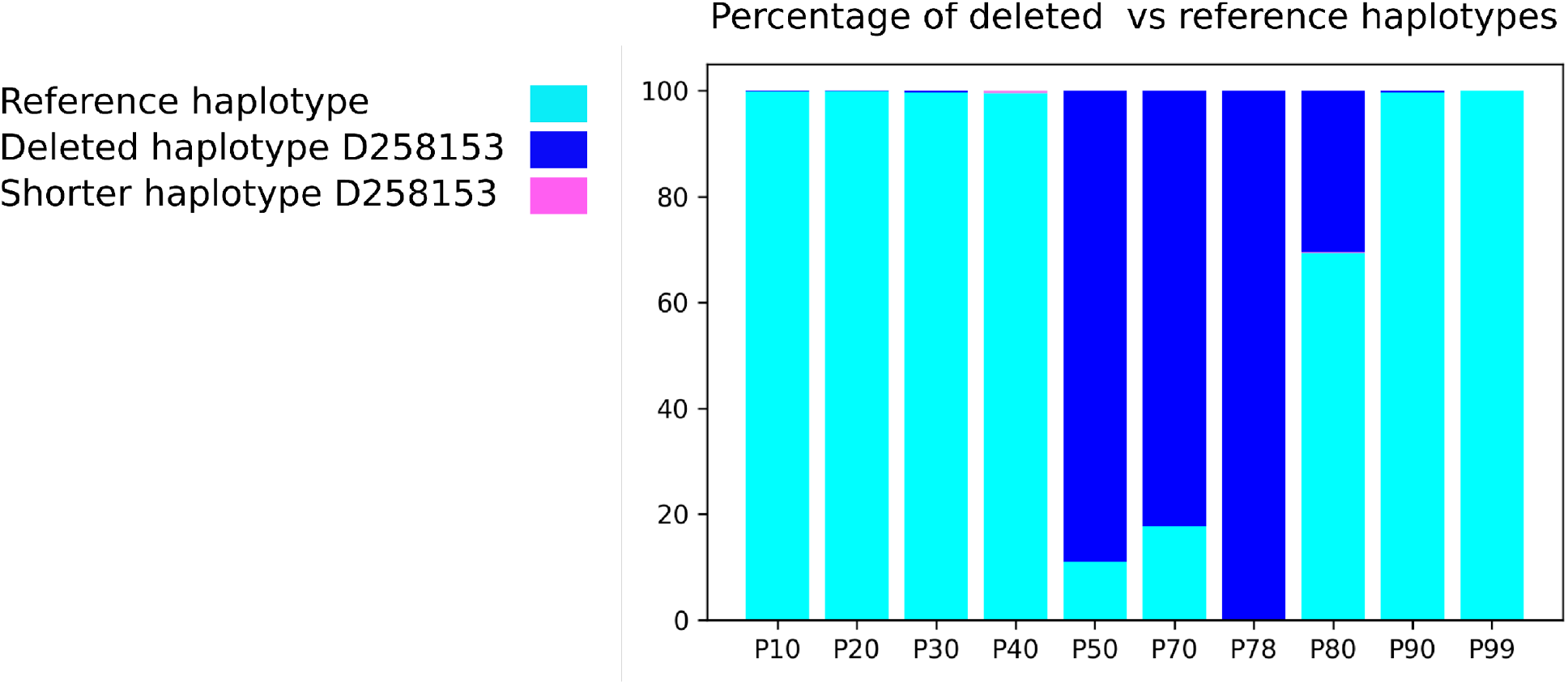
The prevalence of the deleted haplotype during the 10 intermediate passages (P10, P20, P30, P40, P50, P70, P78, P80, P90, P99

This analysis also evidenced several other SVs that alter the structure of ORF150 and of its upstream region, including the beginning of ORF149 (Figure 5). Besides the large deletion, inversions and insertions were also observed in the ORF149-ORF150 region. Inversions were at a low frequency (between 0.01% and 0.53% of the supporting reads) in all passages except for P70, P90 and P99. P10 and P40 showed the lowest and the highest inversion frequencies, respectively (Figure 5, Table S4 (Cherif, 2022)). A large insertion of about 1 kb appeared in P50 and P70 at moderate frequencies (14,34% and 16.01% of the supporting reads, respectively) to disappear in P78 and re-appear at a lower frequency (6,79% of the supporting reads) in P80. The consensus sequence of this insertion corresponds to the fragment 259,517-260,477 of the KHV genome, with an identity of about 90%. In P90, an intriguing inverted-duplicated haplotype was observed at a low proportion (0.054% of the supporting reads). Surprisingly, P99 exhibited a unique reference-like, SV-free haplotype (Figure 5, Table S4 (Cherif, 2022)). All the variations deeply impacted the structure of ORF150 - and sometimes that of ORF149 as well - by shrinking or increasing its size, causing the ORF149 and ORF150 fusion, inverting the ORF150 sequences and duplicating the ORF150 with the deleted, inserted, inverted and inverted-duplicated haplotypes (Figure 5).

**Figure 5:**
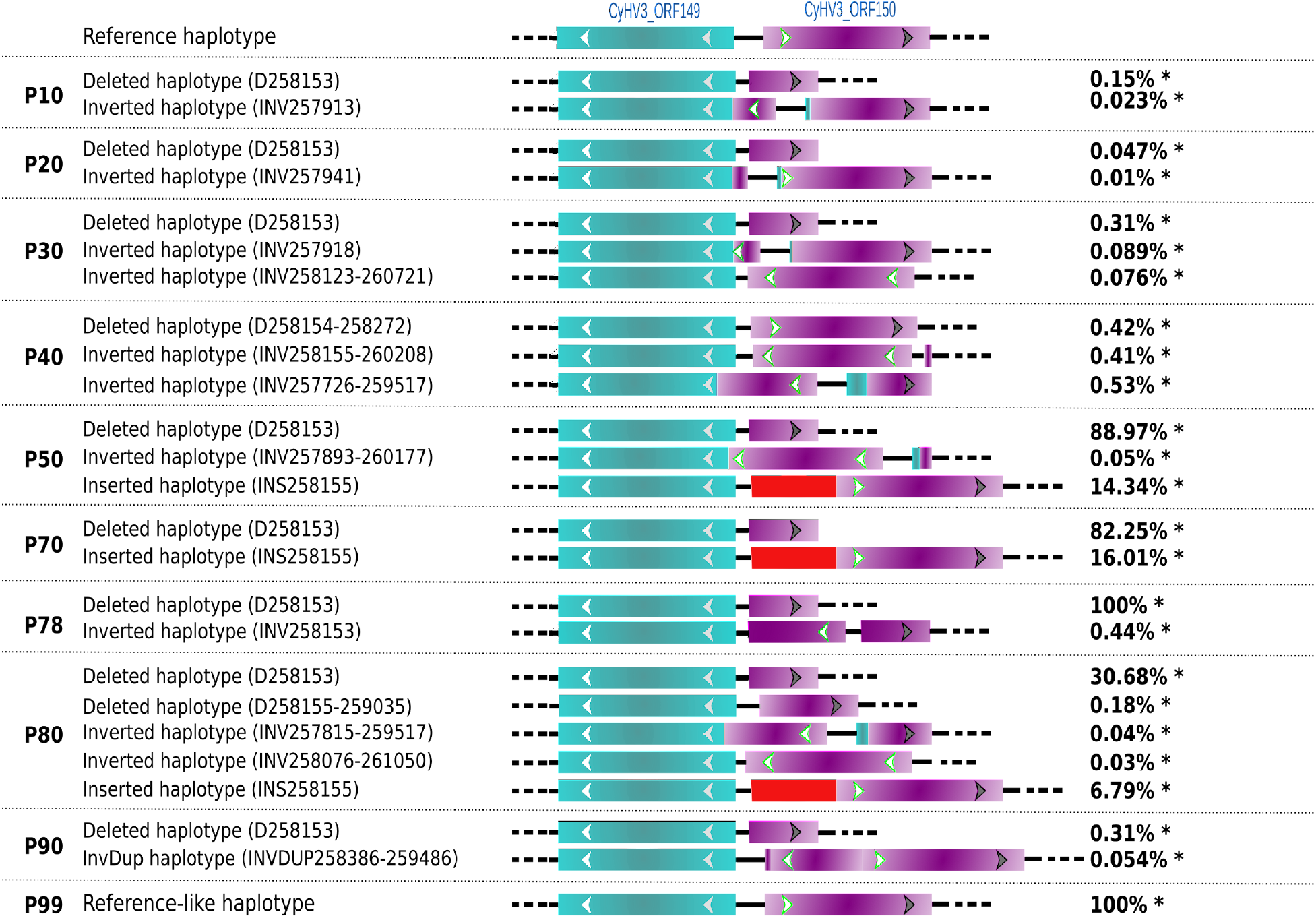
Impact of SV dynamics on the ORF149-ORF150 structure in the successive passages P10, P20, P30, P40, P50, P70, P78, P80, P90 and P99. The inversion is highlighted by an inverted arrow compared to the reference haplotype. Red blocks correspond to an inserted sequence. InvDup = Inverted-duplicated haplotype. *Percentage of reads covering the SVs at given breakpoints.

## Discussion

SVs significantly impact the adaptation of viruses to their natural host and environment (Pérez-Losada et al., 2015). Yet the role of SV diversity and dynamics in large DNA viruses is barely known. Ultra-deep long-read sequencing opens unprecedented ways to gain insights into these untapped viral genome polymorphisms. The present study started to tackle the impact of SVs in the evolution of the large dsDNA KHV during cell culture serial passages using ultra-deep whole-genome and amplicon-based sequencing. The sequence data showed a wide distribution of various SVs along the genome associated with high SV turnover dynamics during the *in vitro* infection cycles and a clear bias towards inversion and deletion events. Analysis of the pathogenicity-associated ORF150 region in ten serial passages mainly highlighted deletions, inversions and insertions that deeply altered the structure of ORF150.

Serial passages of viruses in cell culture may lead to the accumulation of mutations and gene disruptions (Spatz 2010; Colgrove et al., 2014). These mutations can modify viral adaptation and increase or decrease virulence (Boutier et al., 2017; López-Muñoz et al., 2021; Vancsok et al., 2017). In the case of KHV, a previous work using short-read sequencing showed that 99 consecutive *in vitro* passages onto CCB cells resulted in the accumulation of less than 60 small variations (<100 nt) (Klafack et al., 2019). It also showed that the haplotype composition can quickly vary along with infection cycles of KHV *in vitro.* The present study unexpectedly highlighted a high number of structural variations: 87 for P99 and 236 for P78. In contrast, the accumulation of small variations was consistent with what had been observed with short-read sequencing (Klafack et al, 2019). These findings illustrate that long-read sequencing is highly suitable for genome-wide comparisons of viruses. Most importantly, they revealed a hidden source of virus diversification, which had never been reported so far for KHV. They also confirmed that P78 consists of a mixture of undeleted and deleted haplotypes revealing that the undeleted haplotype did not correspond to the native one but to an inverted version of it. We experimentally validated this inversion in P78 by PCR followed by sequencing. Moreover, the inversion found in the middle of the reads excluded the formation of *in silico* chimeras, i.e., chimeras resulting from the basecaller when two molecules are sequenced in the same pore that undergoes fast reloading (Martin and Legget, 2021). These structural variations form a ‘mosaic’ of viral subpopulations that seem to result from multiple rearrangement events, mainly inversions and deletions and, to a lesser extent, insertions and duplications. Such sub-viral variants often lead to Defective Viral Genomes (DVGs) (Vignuzzi and López 2019). Because of their negative impact on viral replication, some forms of DVGs have been extensively studied, and three pathogenesis-related functions have been well-described: interference with viral replication, immunostimulation, and viral persistence (Marriott and Dimmock 2010; Vignuzzi and López 2019). DVGs play a role in viral production interference by accumulating at higher rates than the full-length viral genomes and consequently interfere with viral replication by taking up the polymerase activity and competing for structural proteins (Calain and Roux 1995; Portner and Kingsbury 1971). In addition, DVGs can act as primary stimuli and trigger antiviral immunity by inducing the expression of some interleukins and pro-inflammatory cytokines (for review, see Vignuzzi and López 2019; Xiao et al, 2021). However, the implication of DVGs for virus persistence is complex and still unclear. Nevertheless, a combination of viral replication interfering and cycle asynchrony between full-length viruses and DVGs seems to establish chronic infection (for review, see Vignuzzi and López 2019).

The observed dynamics of structural variations in the KHV genomes during cell culture passages where selection pressures are virtually non-existent may indicate an accumulation of DVGs impacting the pathogenicity. Indeed, Klafack et al. (2019) showed that serial passages significantly attenuated this infectious virus. This attenuation was associated with a truncated KHV genome bearing a 1.3-kb deletion that removes the majority of the ORF150. A very recent study showed that partial or complete depletion of ORF150 leads to a clear attenuation of the virus. It also brought *in vivo* evidence that ORF150 may down-regulate the inflammatory response of carp to enhance viral proliferation, thus confirming a key role of ORF150 product in KHV virulence (Klafack et al., 2022 submitted). ORF150 encodes a protein with a Really Interesting New Gene (RING) domain that is predicted to span 628 amino acids (Li et al, 2015; Aoki et al, 2007). This RING domain contains the HC (C3HC4) type of RING structure, which is involved in the ubiquitination pathway, by acting as an E3 ubiquitin-protein ligase (He and Kwang 2008). Viruses that encode RING finger like-ubiquitin ligases (E3s)may evade host immune responses and also hijack the host’s RING E3 to enhance their replication (Zhang et al.,2018). In aquatic viruses, RING family genes have been reported to be involved in virus latency, replication, and host protein degradation (Shekar and Venugopal 2019; Wang et al., 2021).

As previously observed, the most significant difference affecting all viral haplotypes between P78 and P99 was around the ORF150. For this reason, we focused on this interesting region to determine the passage number at which the presence of the deletion inversion first arose, by sequencing selected representative passages (P10, P20, P30, P40, P50, P70, P80 and P90). We found that deleted and inverted haplotypes appeared as soon as P10 and that their proportion varied along the successive passages. The most striking feature was the rapid and total disappearance of the deletion between P78 and P99, raising many questions regarding the mechanisms that led to the clearance of this major haplotype.

The rapid SV turnover of DNA viruses, including herpesviruses, likely involves recombination (Szpara and VanDoorslaer, 2021; Renner and Szpara, 2018; Cudini et al., 2019; Wilkinson and Weller, 2003; Kolb et al., 2017; Tomer et al., 2019), which is often linked to replication and DNA repair, as well as errors during viral genome replication (Kulkarni and Fortunato 2011; Xiaofei and Kowalik 2014). In the present case, the good conservation of breakpoints around ORF150 may signify homologous recombination. However, whether this recombination occurs within the same genomic entities or between different viruses remains open. It would be interesting to assess the involvement of each of these mechanisms in generating the observed structural diversity. The multiple mechanisms of DNA virus evolution beyond single nucleotide substitutions likely confer KHV a high level of evolutionary adaptability.

Classically, the generation of live-attenuated vaccines is achieved by passaging the virus in cell culture under different conditions (in different host species or at lower or higher temperatures), in order to induce mutation accumulation that supports viral adaptation to the specific conditions and provides viral attenuation (Minor, 2015, Hanley 2011). With the exception of the OPV polio vaccine viruses (Kew et al., 2005), the exact mechanisms by which these mutations lead to attenuated phenotypes are usually poorly characterized (Lauring et al., 2010). However, live-attenuated viruses can revert to virulent phenotypes either by reversions (as shown here between P78 and P99), the introduction of compensatory mutations, or recombination with viruses belonging to the same genus (Cann et al., 1984; Bull et al., 2018; Muslin et al., 2019). Additionally, the combination of multiple live-attenuated viruses may result in competition or facilitation between individual vaccine viruses, resulting in undesirable increases in virulence or decreases in immunogenicity (Hanley 2011; Pereira-Gomez et al., 2021). Recently, genetic engineering has led to many novel approaches to generating live-attenuated virus vaccines containing modifications to prevent virulence reversion (Yeh et al., 2020) and improve interferences among multiple vaccine strains (Pereira-Gomez et al., 2021).

## Conclusion

Our findings confirm that CyHV-3 can evolve rapidly during infectious cycles in cell culture, and SVs are a major component in the evolutionary process of this virus. SVs are extremely dynamic under *in vitro* controlled conditions, and it would now be interesting to evaluate their dynamics *in vivo.* The present study also contributes to the basic research on the mechanisms underlying attenuation and may have important outcomes for the design of safe live-attenuated vaccine formulations.

## Supporting information

Supplementary_Information_Fuandila_etal

## Acknowledgments

Part of this work was supported by the “Laboratoire d’excellence” (LabEx) CeMEB, an ANR “Investissements d’avenir” program (ANR-10-LABX-04-01), through the exploratory research project HaploFit. This work benefited from the Montpellier Bioinformatics Biodiversity platform supported by the LabEx CeMEB, an ANR “Investissements d’avenir” program (ANR-10-LABX-04-01). Real-time PCR data were produced on the qPHD platform of Montpellier University, with the support of LabEx CeMEB. N. N. F benefited from a CampusFrance fellowship for her PhD thesis, and we are very grateful to the French Embassy of Indonesia. E. C received a post-doctoral fellowship from IRD. Version 4 of this preprint has been peer-reviewed and recommended by Peer Community In Infections (https://doi.org/10.24072/pci.infections.100001).

## Data, scripts and codes availability

Raw sequences (fastq files) were stored in the public Sequence Read Archive (SRA) repository and can be accessed under the bioproject PRJNA511566. The original genetic data yielded by genotyping (in vcf format), codes and additional metadata are available in a publicly-available OSF repository: https://doi.org/10.17605/osf.io/3c2ag.

## Supplementary material

Supplementary material is available online: https://doi.org/10.17605/osf.io/3c2ag.

## Authors’ contributions

**Nurul Novelia Fuandila:** Investigation; Methodology; Formal analysis; Writing – original draft; Data Validation

**Anne-Sophie Gosselin-Grenet:** Investigation; Methodology; Data curation; Data Validation; Writing – original draft

**Marie-Ka Tilak:** Investigation; Methodology; Data Validation

**Sven M Bergmann**: Resources; Writing – review and editing

**Sandro Klafack:** Resources; Writing – review and editing

**Angela Lusiastuti:** Methodology; Writing – review and editing

**Munti Yuhanna:** Methodology; Writing – review and editing

**Jean-Michel Escoubas:** Methodology; Funding acquisition; Writing – review and editing

**Anna-Sophie Fiston-Lavier:** Writing – review and editing

**Jean-Christophe Avarre:** Conceptualization; Data curation; Methodology; Funding acquisition; Data Validation; Writing – original draft

**Emira Cherif:** Conceptualization; Data curation; Formal analysis; Software; Writing – original draft;

## References

Akhtar, L. N., Bowen, C. D., Renner, D. W., Pandey, U., Della Fera, A. N., Kimberlin, D. W.,… & Szpara, M. L. (2019). Genotypic and phenotypic diversity of herpes simplex virus 2 within the infected neonatal population. MSphere, 4(1), e00590–18. https://doi.org/10.1128/mSphere.00590-18

Aoki, T., Hirono, I., Kurokawa, K., Fukuda, H., Nahary, R., Eldar, A., … & Hedrick, R. P. (2007). Genome sequences of three koi herpesvirus isolates representing the expanding distribution of an emerging disease threatening koi and common carp worldwide. Journal of virology, 81(10), 5058–5065. https://doi.org/10.1128/JVI.00146-07

Boutier, M., Ronsmans, M., Rakus, K., Jazowiecka-Rakus, J., Vancsok, C., Morvan, L., … & Vanderplasschen, A. (2015). Cyprinid herpesvirus 3: an archetype of fish alloherpesviruses. Advances in Virus Research, 93, 161–256. https://doi.org/10.1016/bs.aivir.2015.03.001

Boutier, Maxime; Gao, Yuan; Vancsok, Catherine; Suárez, Nicolás M.; Davison, Andrew J.; Vanderplasschen, Alain (2017): Identification of an essential virulence gene of cyprinid herpesvirus 3. In: Antiviral research 145, S. 60–69. https://doi.org/10.1016/j.antiviral.2017.07.002

Bull, J. C., Godfray, H. C. J., & O’Reilly, D. R. (2003). A few-polyhedra mutant and wild-type nucleopolyhedrovirus remain as a stable polymorphism during serial coinfection in Trichoplusia ni. Applied and environmental microbiology, 69(4), 2052–2057. https://doi.org/10.1128/AEM.69.4.2052-2057.2003

Bull, J. J., Smithson, M. W., & Nuismer, S. L. (2018). Transmissible viral vaccines. Trends in microbiology, 26(1), 6–15. https://doi.org/10.1016/j.tim.2017.09.007

Calain, P., & Roux, L. Functional characterisation of the genomic and antigenomic promoters of Sendai virus. Virology 212, 163–173 (1995). https://doi.org/10.1006/viro.1995.1464

Cann, A. J., Stanway, G., Hughes, P. J., Minor, P. D., Evans, D. M. A., Schild, G. T., & Almond, J. (1984). Reversion to neurovirulence of the live-attenuated Sabin type 3 oral pollovirus vaccine. Nucleic acids research, 12(20), 7787–7792. https://doi.org/10.1093/nar/12.20.7787

Chateigner, A., Bézier, A., Labrousse, C., Jiolle, D., Barbe, V., & Herniou, E. A. (2015). Ultra deep sequencing of a baculovirus population reveals widespread genomic variations. Viruses, 7(7), 3625–3646. https://doi.org/10.3390/v7072788

Cherif, E. (2022). KHV_SVProject. OSF, https://doi.org/10.17605/OSF.IO/3C2AG

Colgrove, R., Diaz, F., Newman, R., Saif, S., Shea, T., Young, S., … & Knipe, D. M. (2014). Genomic sequences of a low passage herpes simplex virus 2 clinical isolate and its plaque-purified derivative strain. Virology, 450, 140–145. https://doi.org/10.1016/j.virol.2013.12.014

Cudini, J., Roy, S., Houldcroft, C. J., Bryant, J. M., Depledge, D. P., Tutill, H., … & Breuer, J. (2019). Human cytomegalovirus haplotype reconstruction reveals high diversity due to superinfection and evidence of within-host recombination. Proceedings of the National Academy of Sciences, 116(12), 5693–5698. https://doi.org/10.1073/pnas.1818130116

Fuchs, Walter; Granzow, Harald; Dauber, Malte; Fichtner, Dieter; Mettenleiter, Thomas C. (2014): Identification of structural proteins of koi herpesvirus. In: Archives of virology 159 (12), S.3257–3268. https://doi.org/10.1007/s00705-014-2190-4

Gao, Y., Suárez, N. M., Wilkie, G. S., Dong, C., Bergmann, S., Lee, P. Y. A.,… & Boutier, M. (2018). Genomic and biologic comparisons of cyprinid herpesvirus 3 strains. Veterinary research, 49(1), 1–11. https://doi.org/10.1186/s13567-018-0532-z

Gotesman, M., Kattlun, J., Bergmann, S. M., & El-Matbouli, M. (2013). CyHV-3: the third cyprinid herpesvirus. Diseases of aquatic organisms, 105(2), 163–174. https://doi.org/10.3354/dao02614

Haenen, O. L. M., Way, K., Bergmann, S. M., & Ariel, E. (2004). The emergence of koi herpesvirus and its significance to European aquaculture. Bulletin of the European Association of Fish Pathologists, 24(6), 293–307.

Hammoumi, Saliha; Vallaeys, Tatiana; Santika, Ayi; Leleux, Philippe; Borzym, Ewa; Klopp, Christophe; Avarre, Jean-Christophe (2016): Targeted genomic enrichment and sequencing of CyHV-3 from carp tissues confirms low nucleotide diversity and mixed genotype infections. In: PeerJ, e2516. https://doi.org/10.7717/peerj.2516

Hanley, K. A. (2011). The double-edged sword: How evolution can make or break a live-attenuated virus vaccine. Evolution: Education and Outreach, 4(4), 635–643. https://doi.org/10.1007/s12052-011-0365-y

He, F., & Kwang, J. (2008). Identification and characterization of a new E3 ubiquitin ligase in white spot syndrome virus involved in virus latency. Virology journal, 5, 151. https://doi.org/10.1186/1743-422X-5-151

Hedrick, R. P., Gilad, O., Yun, S., Spangenberg, J. V., Marty, G. D., Nordhausen, R. W., … & Eldar, A. (2000). A herpesvirus associated with mass mortality of juvenile and adult koi, a strain of common carp. Journal of Aquatic Animal Health, 12(1), 44–57. https://doi.org/10.1007/s12052-011-0365-y

Jeffares, D. C., Jolly, C., Hoti, M., Speed, D., Shaw, L., Rallis, C., … & Sedlazeck, F. J. (2017). Transient structural variations have strong effects on quantitative traits and reproductive isolation in fission yeast. Nature communications, 8(1), 1–11. https://doi.org/10.1038/NCOMMS14061

Kew, O. M., Sutter, R. W., de Gourville, E. M., Dowdle, W. R., & Pallansch, M. A. (2005). Vaccine-derived polioviruses and the endgame strategy for global polio eradication. Annu. Rev. Microbiol., 59, 587–635. https://doi.org/10.1146/annurev.micro.58.030603.123625

Klafack, S., Fiston-Lavier, A. S., Bergmann, S. M., Hammoumi, S., Schröder, L., Fuchs, W.,… & Avarre, J. C. (2019). Cyprinid herpesvirus 3 evolves in vitro through an assemblage of haplotypes that alternatively become dominant or under-represented. Viruses, 11(8), 754. https://doi.org/10.3390/v11080754

Klafack, S., Schröder, L., Jin, Y., Lenk, M., Lee, P.Y., Fuchs, W, Avarre, J. C., Bergmann, S. M. (2022). Development of an attenuated vaccine against Koi Herpesvirus Disease (KHVD) suitable for oral administration and immersion. (submitted, NPJ Vaccines)

Klafack, S., Wang, Q., Zeng, W., Wang, Y., Li, Y., Zheng, S.C., Kempter, J., Lee, P.Y., & Bergmann, S. M. (2017). Genetic Variability of Koi Herpesvirus In vitro—A Natural Event?. Front. Microbiol., 8, 982. https://doi.org/10.3389/fmicb.2017.00982

Kolb, A. W., Lewin, A. C., Moeller Trane, R., McLellan, G. J., & Brandt, C. R. (2017). Phylogenetic and recombination analysis of the herpesvirus genus varicellovirus. BMC genomics, 18(1), 1–17. https://doi.org/10.1186/s12864-017-4283-4

Kulkarni, A. S., & Fortunato, E. A. (2011). Stimulation of homology-directed repair at I-SceI-induced DNA breaks during the permissive life cycle of human cytomegalovirus. Journal of virology, 85(12), 6049–6054. https://doi.org/10.1128/JVI.02514-10

Lauring, A. S., & Andino, R. (2010). Quasispecies theory and the behavior of RNA viruses. PLoS pathogens, 6(7), e1001005. https://doi.org/10.1371/journal.ppat.1001005

Lauring, A. S., Jones, J. O., & Andino, R. (2010). Rationalizing the development of live attenuated virus vaccines. Nature biotechnology, 28(6), 573–579. https://doi.org/10.1038/nbt.1635

Lecompte, L., Peterlongo, P., Lavenier, D., & Lemaitre, C. (2020). SVJedi: Genotyping structural variations with long reads. Bioinformatics, 36(17), 4568–4575. https://doi.org/10.1093/bioinformatics/btaa527

Li, D., Lott, W. B., Lowry, K., Jones, A., Thu, H. M., & Aaskov, J. (2011). Defective interfering viral particles in acute dengue infections. PloS one, 6(4), e19447. https://doi.org/10.1371/journal.pone.0019447

Li, H. (2018). Minimap2: pairwise alignment for nucleotide sequences. Bioinformatics, 34(18), 3094–3100. https://doi.org/10.1093/bioinformatics/bty191

Li, H., Handsaker, B., Wysoker, A., Fennell, T., Ruan, J., Homer, N., … & Durbin, R. (2009). The sequence alignment/map format and SAMtools. Bioinformatics, 25(16), 2078–2079. https://doi.org/10.1093/bioinformatics/btp352

Li, W., Lee, X., Weng, S., He, J., Dong, C. (2015). Whole-genome sequence of a novel Chinese cyprinid herpesvirus 3 isolate reveals the existence of a distinct European genotype in East Asia. Veterinary microbiology 175 (2-4), 185–194. https://doi.org/10.1016/j.vetmic.2014.11.022

Loiseau, V., Herniou, E. A., Moreau, Y., Lévêque, N., Meignin, C., Daeffler, L.,… & Gilbert, C. (2020). Wide spectrum and high frequency of genomic structural variation, including transposable elements, in large double-stranded DNA viruses. Virus evolution, 6(1), vez060. https://doi.org/10.1093/ve/vez060

López-Muñoz, A. D., Rastrojo, A., Martín, R., & Alcamí, A. (2021). Herpes simplex virus 2 (HSV-2) evolves faster in cell culture than HSV-1 by generating greater genetic diversity. PLoS pathogens, 17(8), e1009541. https://doi.org/10.1371/journal.ppat.1009541

Marriott, A. C., and Dimmock, N. J. (2010) ‘Defective Interfering Viruses and Their Potential as Antiviral Agents’, Reviews in Medical Virology, 20: 51–62. https://doi.org/10.1002/rmv.641

Martin, S., & Leggett, R. M. (2021). Alvis: a tool for contig and read ALignment VISualisation and chimera detection. BMC bioinformatics, 22(1), 1–10. https://doi.org/10.1186/s12859-021-04056-0

Minor, P. D. (2015). Live attenuated vaccines: Historical successes and current challenges. Virology, 479, 379–392. https://doi.org/10.1016/j.virol.2015.03.032

Molenkamp, R., Rozier, B. C., Greve, S., Spaan, W. J., & Snijder, E. J. (2000). Isolation and characterization of an arterivirus defective interfering RNA genome. Journal of virology, 74(7), 3156–3165. https://doi.org/10.1128/JVI.74.7.3156-3165.2000

Muslin, C., Mac Kain, A., Bessaud, M., Blondel, B., & Delpeyroux, F. (2019). Recombination in enteroviruses, a multi-step modular evolutionary process. Viruses, 11(9), 859. https://doi.org/10.3390/v11090859

O’Hara, P. J., Nichol, S. T., Horodyski, F. M., & Holland, J. J. (1984). Vesicular stomatitis virus defective interfering particles can contain extensive genomic sequence rearrangements and base substitutions. Cell, 36(4), 915–924. https://doi.org/10.1016/0092-8674(84)90041-2

Pereira-Gómez, M., Carrau, L., Fajardo, Á., Moreno, P., & Moratorio, G. (2021). Altering Compositional Properties of Viral Genomes to Design Live-Attenuated Vaccines. Frontiers in Microbiology, 1706. https://doi.org/10.3389/fmicb.2021.676582

Pérez-Losada, M., Arenas, M., Galán, J. C., Palero, F., & González-Candelas, F. (2015). Recombination in viruses: mechanisms, methods of study, and evolutionary consequences. Infection, Genetics and Evolution, 30, 296–307. https://doi.org/10.1016/j.meegid.2014.12.022

Portner, A., & Kingsbury, D. W. Homologous interference by incomplete Sendai virus particles: changes in virus-specifc ribonucleic acid synthesis. J. Virol. 8, 388–394 (1971). https://doi.org/10.1128/jvi.8.4.388-394.1971

Rakus, K., Ouyang, P., Boutier, M., Ronsmans, M., Reschner, A., Vancsok, C., … & Vanderplasschen, A. (2013). Cyprinid herpesvirus 3: an interesting virus for applied and fundamental research. Veterinary Research, 44(1), 1–16. https://doi.org/10.1186/1297-9716-44-85

Ramírez, F., Ryan, D. P., Grüning, B., Bhardwaj, V., Kilpert, F., Richter, A. S., … & Manke, T. (2016). deepTools2: a next generation web server for deep-sequencing data analysis. Nucleic acids research, 44(W1), W160–W165. https://doi.org/10.1093/nar/gkw257

Renner, D. W., & Szpara, M. L. (2018). Impacts of genome-wide analyses on our understanding of human herpesvirus diversity and evolution. Journal of virology, 92(1), e00908–17. https://doi.org/10.1128/JVI.00908-17

Renzette, N., Pokalyuk, C., Gibson, L., Bhattacharjee, B., Schleiss, M. R., Hamprecht, K., … & Kowalik, T. F. (2015). Limits and patterns of cytomegalovirus genomic diversity in humans. Proceedings of the National Academy of Sciences, 112(30), E4120–E4128. https://doi.org/10.1073/pnas.1501880112

Sanjuán, R., & Domingo-Calap, P. (2016). Mechanisms of viral mutation. Cellular and molecular life sciences, 73(23), 4433–4448. https://doi.org/10.1007/s00018-016-2299-6

Sedlazeck, F. J., Rescheneder, P., Smolka, M., Fang, H., Nattestad, M., Von Haeseler, A., & Schatz, M. C. (2018). Accurate detection of complex structural variations using single-molecule sequencing. Nature methods, 15(6), 461–468. doi: 10.1038/s41592-018-0001-7

Shekar, M., & Venugopal, M. N. (2019). Identification and characterization of novel double zinc fingers encoded by putative proteins in genome of white spot syndrome virus. Archives of virology, 164(4), 961–969. https://doi.org/10.1007/s00705-019-04150-y

Spatz, S. J. (2010). Accumulation of attenuating mutations in varying proportions within a high passage very virulent plus strain of Gallid herpesvirus type 2. Virus research, 149(2), 135–142. https://doi.org/10.1016/j.virusres.2020.01.007

Sunarto, A., McColl, K. A., Crane, M. S. J., Sumiati, T., Hyatt, A. D., Barnes, A. C., & Walker, P. J. (2011). Isolation and characterization of koi herpesvirus (KHV) from Indonesia: identification of a new genetic lineage. Journal of fish diseases, 34(2), 87–101. https://doi.org/10.1111/j.1365-2761.2010.01216.x

Szpara, M.L., & Van Doorslaer, K. Mechanisms of dna virus evolution. In Dennis H. Bamford and Mark Zuckerman, editors, Encyclopedia of Virology (Fourth Edition),pages 71–78. Academic Press, Oxford, fourth edition edition, 2021. 3 https://doi.org/10.1016/B978-0-12-809633-8.20993-X

Te Yeh, M., Bujaki, E., Dolan, P. T., Smith, M., Wahid, R., Konz, J., … & Andino, R. (2020). Engineering the live-attenuated polio vaccine to prevent reversion to virulence. Cell host & microbe, 27(5), 736–751. https://doi.org/10.1016/j.chom.2020.04.003

Tomer, E., Cohen, E. M., Drayman, N., Afriat, A., Weitzman, M. D., Zaritsky, A., & Kobiler, O. (2019). Coalescing replication compartments provide the opportunity for recombination between coinfecting herpesviruses. The FASEB journal, 33(8), 9388–9403. https://doi.org/10.1096/fj.201900032R

Vancsok, C., Peñaranda, M. M. D., Raj, V. S., Leroy, B., Jazowiecka-Rakus, J., Boutier, M., … & Vanderplasschen, A. F. (2017). Proteomic and functional analyses of the virion transmembrane proteome of cyprinid herpesvirus 3. Journal of Virology, 91(21), e01209–17. DOI: 10.1128/JVI.01209-17

Vignuzzi, M., & López, C. B. (2019). Defective viral genomes are key drivers of the virus–host interaction. Nature microbiology, 4(7), 1075–1087. https://doi.org/10.1038/s41564-019-0465-y

Wang, Z. H., Ke, F., Zhang, Q. Y., & Gui, J. F. (2021). Structural and functional diversity among five ring finger proteins from Carassius auratus herpesvirus (CaHV). Viruses, 13(2), 254. https://doi.org/10.3390/v13020254

White, R., Pellefigues, C., Ronchese, F., Lamiable, O., & Eccles, D. (2017). Investigation of chimeric reads using the MinION. F1000Research, 6. https://doi.org/10.12688/f1000research.11547.2

Wilkinson, D., & Weller, S. (2003). The role of DNA recombination in herpes simplex virus DNA replication. IUBMB life, 55(8), 451–458. https://doi.org/10.1080/15216540310001612237

Wilkinson, Dianna E., Weller, Sandra K. (2003): The role of DNA recombination in herpes simplex virus DNA replication. In: IUBMB life 55 (8), S. 451–458. https://doi.org/10.1080/15216540310001612237

Xiao, Y., Lidsky, P. V., Shirogane, Y., Aviner, R., Wu, C. T., Li, W., … & Andino, R. (2021). A defective viral genome strategy elicits broad protective immunity against respiratory viruses. Cell, 184(25), 6037–6051. https://doi.org/10.1016/j.cell.2021.11.02

Xiaofei, E., & Kowalik, T.F. The dna damage response induced by infection with human cytomegalovirus and other viruses. Viruses, 6(5): 2155–2185, 2014. https://doi.org/10.3390/v6052155

Zhang, Y., Li, L.F., Munir, M. Qiu, H. J. (2018). RING-Domain E3 Ligase-Mediated Host-Virus Interactions: Orchestrating Immune Responses by the Host and Antagonizing Immune Defense by Viruses. Frontiers in immunology, 9, 1083. https://doi.org/10.3389/fimmu.2018.01083

