## Supplementary_Information_Fuandila_etal for "Structural variation turnovers and defective genomes: key drivers for the in vitro evolution of the large double-stranded DNA koi herpesvirus (KHV)"

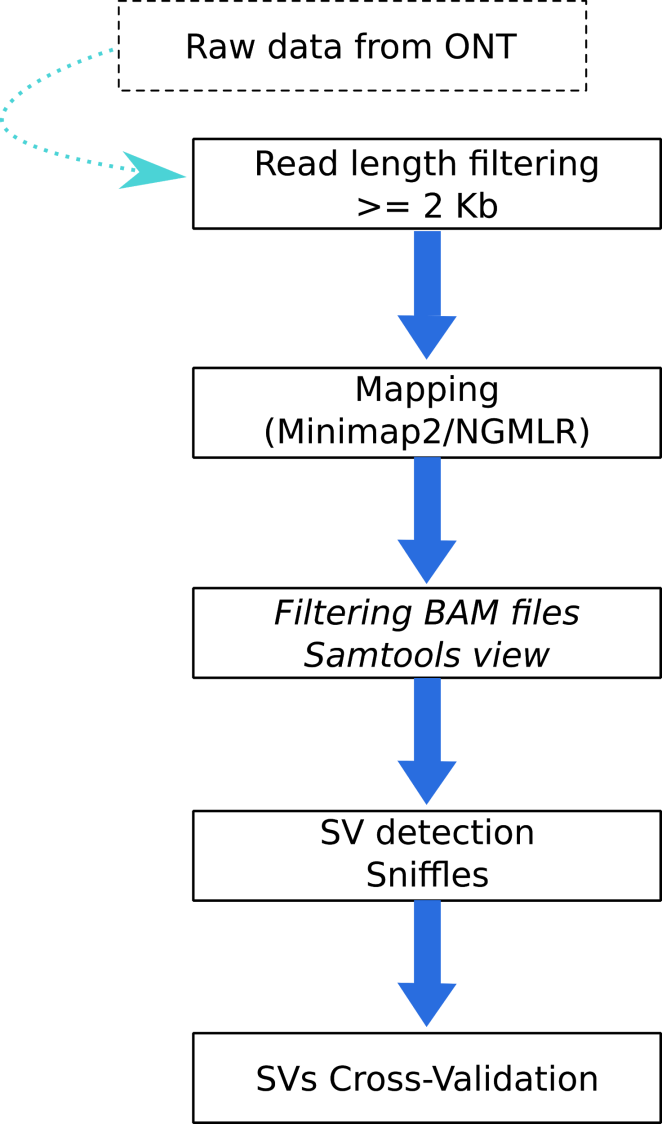

**Figure S1.** Overview of the bioinformatic workflow to detect SVs in P78 and P99 genomes.

**Table S1** Main features of the obtained raw reads for each strain.

num_seq= number of reads, min_len= minimum length, avg_len= average length, max_len= maximum length

| **Sample** | **num_seqs** | **min_len** | **avg_len** | **max_len** | **N50** | **Q20(%)** | **Q30(%)** |
| --- | --- | --- | --- | --- | --- | --- | --- |
| P78_all_fast5_pass_basecalled_gup4.fastq | 4900000 | 25 | 1080.7 | 91494 | 1598 | 59.92 | 18.65 |
| P99_all_fast5_pass_basecalled_gup4.fastq | 2293830 | 96 | 3764.5 | 279443 | 7643 | 57.25 | 17.94 |
| P10_all.fastq | 244000 | 91 | 1144.4 | 7819 | 2022 | 45.31 | 14.70 |
| P20_all.fastq | 484000 | 96 | 1091.9 | 7146 | 1938 | 45.44 | 14.74 |
| P30_all.fastq | 472000 | 85 | 1455.1 | 7519 | 2430 | 45.40 | 14.62 |
| P40_all.fastq | 612000 | 99 | 1003.4 | 6197 | 1635 | 44.62 | 14.33 |
| P50_all.fastq | 115135 | 110 | 1068.3 | 4596 | 1632 | 37.68 | 9.88 |
| P70_all.fastq | 65062 | 104 | 1057.3 | 4824 | 1614 | 37.53 | 9.90 |
| P80_all.fastq | 135441 | 99 | 1191.7 | 4592 | 1742 | 37.79 | 9.81 |
| P90_all.fastq | 95635 | 123 | 1502.9 | 7823 | 2624 | 38.18 | 10.10 |

**Table S2** SVs detected by each mapper/caller combination for P78 and P99 strains

|  | **P78** | | **P99** | |
| --- | --- | --- | --- | --- |
|  | **Minimap2+Sniffles** | **NGMLR+Sniffles** | **Minimap2+Sniffles** | **NGMLR+Sniffles** |
| Deletion | 135 | 125 | 22 | 147 |
| Duplication | 49 | 19 | 2 | 30 |
| Inversion | 538 | 308 | 176 | 210 |
| Insertion | 9 | 8 | 10 | 10 |
| Total | **731** | **460** | **210*** | **397*** |

* P99 seems to accumulate complex SVs as Inversion-Duplications, which explains why the combination Minimap2+Sniffles detects fewer SVs than NGMLR+Sniffles while it is the opposite in P78 SVs are mainly Inversions and deletion. (see Sedlazeck et al. 2018 for more details on the mapping/detection algorithms).

**Table S3** ORFs included within SVs distribution windows for P78 and P99. Only SVs supported by more than 0,1 % of the reads were considered.

Abbreviation : Perc_Svs_reads= percentage of reads covering the SV ; AF= Allelic frequence

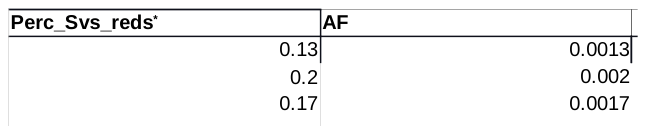

**Table S4** SVs prevalence and type for P10, P20, P30, P40, P50, P70, P78, P80, P90 and P99.

| Strain | Supporting_reads_SVs^*^ | Perc_SVs_reds^*^ | SVs | Ref_  reads^*^ | Perc_ref_  reads^*^ | Total_  reads^*^ | Breakpoints |
| --- | --- | --- | --- | --- | --- | --- | --- |
| P10 | 102 | 0.15 | Del | 66001 | 99.84 | 66103 | 258153-259517 |
| P10 | 13 | 0.023 | Inv | 56244 | 99.97 | 56244 | 257913-258821 |
| P20 | 58 | 0.047 | Del | 121869 | 99.97 | 56257 | 258153-259517 |
| P20 | 10 | 0.01 | Inv | 93483 | 99.99 | 93493 | 257941-258545 |
| P30 | 115 | 0.31 | Del | 36858 | 99.68 | 36973 | 258153-259517 |
| P30 | 33 | 0.089 | Inv | 36858 | 99.91 | 36891 | 257918-258643 |
| P30 | 30 | 0.076 | Inv | 39293 | 99.92 | 39323 | 258123-260721 |
| P40 | 39 | 0.42 | Del | 9121 | 99.57 | 9160 | 258154-258272 |
| P40 | 38 | 0.41 | Inv | 9114 | 99.58 | 9152 | 257726-259517 |
| P40 | 47 | 0.53 | Inv | 8728 | 99.46 | 8775 | 258155-260208 |
| P50 | 7468 | 88.97 | Del | 925 | 11.02 | 8393 | 258153-259517 |
| P50 | 16 | 0.05 | Inv | 31749 | 99.94 | 31765 | 257893-260117 |
| P50 | 525 | 14.34 | Ins | 3136 | 85.65 | 3661 | 258155 |
| P70 | 4038 | 82.25 | Del | 871 | 17.74 | 4909 | 258153-259517 |
| P70 | 322 | 16.01 | Ins | 1688 | 83.98 | 2010 | 258155 |
| P78 | 6902 | 100 | Del | 0 | 0 | 6902 | 258154-259517 |
| P78 | 30 | 0.44 | Inv | 6704 | 99.55 | 6734 | 258155-259518 |
| P80 | 10838 | 30.68 | Del | 24487 | 69.31 | 35325 | 258153-259517 |
| P80 | 21 | 0.18 | Del | 11196 | 99.81 | 11217 | 258155-259035 |
| P80 | 18 | 0.04 | Inv | 36044 | 99.95 | 36062 | 257815-259517 |
| P80 | 12 | 0.03 | Inv | 34628 | 99.96 | 34640 | 258076-261050 |
| P80 | 761 | 6.79 | Ins | 10445 | 93.2 | 11206 | 258155 |
| P90 | 88 | 0.31 | Del | 27560 | 99.68 | 27646 | 258153-259517 |
| P90 | 15 | 0.054 | InvDup | 27273 | 99.94 | 27288 | 258386-259486 |
| P99^+^ | - | 0 |  | - | 100 | - | - |

*At given breakpoints:

Supporting_reads_SVs and Perc_SVs_reds correspond to the number of reads covering a given SV and their percentage.

Ref_reads and Perc_ref_reads correspond to the number of references like reads and their percentage.

Total_reads correspond to the total number of reads covering the breakpoints region.

+As no SVs were detected within the targeted region in P99, 100 % of the mapped reads are reference-like.

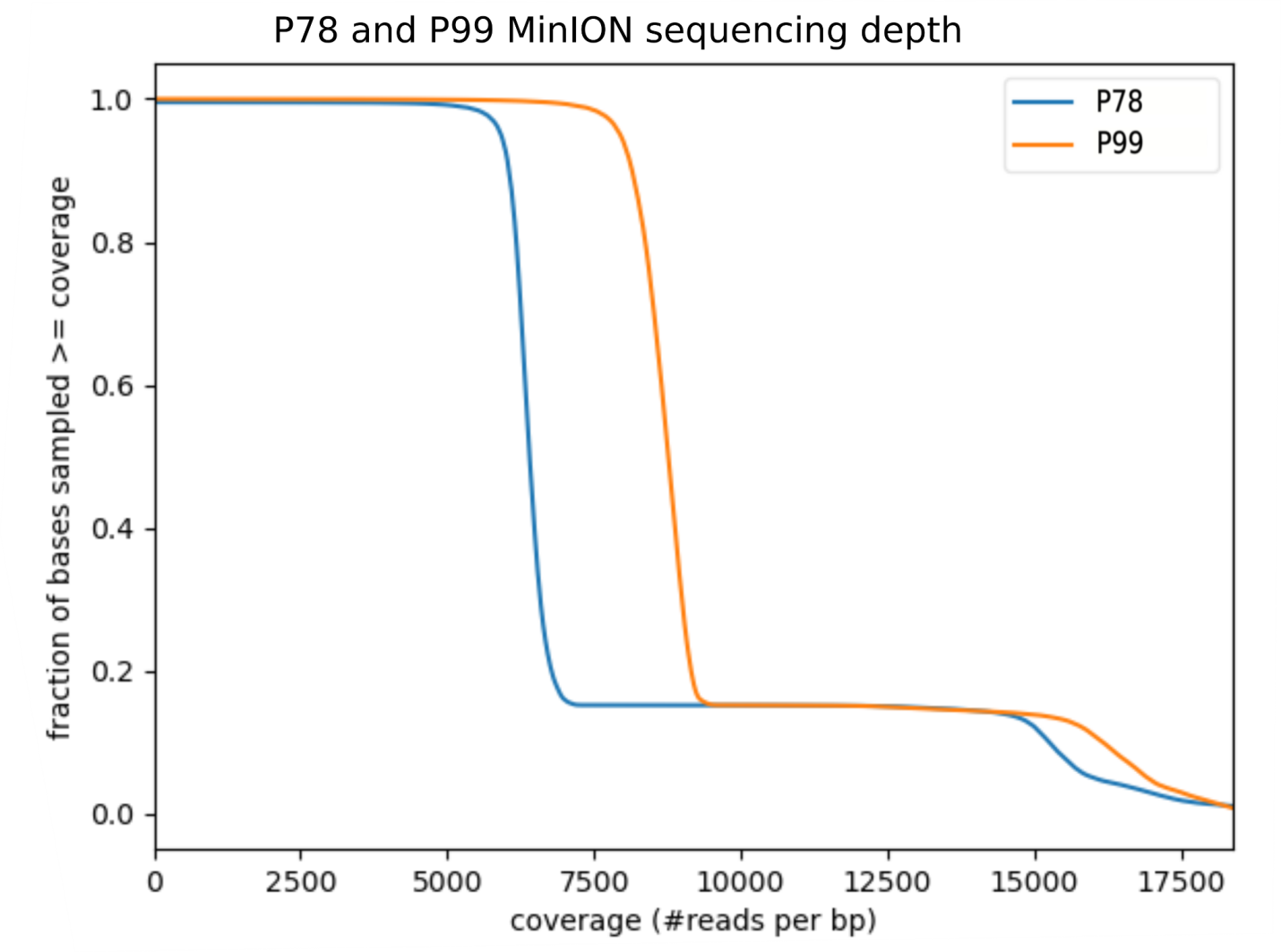

**Figure S2.** Sequencing depth for KHVP78 and KHVP99 samples. This graph shows how the bases are covered and how many times. For example, 100% of the sampled bp from the KHVP99 genome have at least 7500 overlapping reads and around 10% have at least 15000 overlapping reads.
